# Genomically and ecologically cohesive subspecies as biologically meaningful units in the global ocean microbiome

**DOI:** 10.64898/2026.09.29.755032

**Authors:** Lucas Paoli, Taylor Priest, Adrián A. Davín, Melanie L. Stäubli, Dominic Eriksson, Hans-Joachim Ruscheweyh, Martin F. Polz, Shinichi Sunagawa

**Affiliations:** Department of Microbiology, École Polytechnique Fédérale de Lausanne, Lausanne, 1015, Switzerland; Department of Biology, Institute of Microbiology and Swiss Institute of Bioinformatics, ETH Zürich, Zürich, 8092, Switzerland; Department of Environmental Systems Science, Institute of Biogeochemistry and Pollutant Dynamics, ETH Zürich, Zürich, 8092, Switzerland; Division of Microbial Ecology, Centre for Microbiology and Environmental Systems Science, University of Vienna, Vienna, Austria

**Author notes:** These authors have contributed equally to this work (Lucas Paoli, Taylor Priest).

## Abstract

Microbial diversity is organised into operationally defined species. However, it remains unclear whether these, or finer-scale groups represent biologically meaningful units in natural communities. Here, we implement a scalable, cultivation-independent framework to identify 724 genomically discrete and ecologically cohesive subspecies across 270 species in the global ocean microbiome. Their diversification is underpinned by gene-specific, nucleotide-level selection on the core genomic backbone and by variable restructuring of gene content, both of which proceed in a subspecies-specific manner. In over half of the species tested, subspecies were significantly associated with distinct environmental conditions, most frequently temperature. This coupling of genomic discreteness and ecological cohesiveness positions these subspecies as biologically meaningful units, a fundamental level at which ecological and evolutionary processes jointly drive divergence and structure microbial diversity in the oceans.

## INTRODUCTION

Microbial diversity is the result of ecological and evolutionary processes: mutations generate new genotypes, gene flow connects or isolates them, selection sorts them along ecological gradients, and drift acts on them in small or fragmented populations^1–4^. Together, these processes determine which organisms occupy an ecological niche, how biological diversity partitions within and between lineages, and what functions communities perform^5^. These processes act on populations, sets of interconnected individuals from the same species that co-occur in space and time^5–7^. However, microbial populations are difficult to delimit in natural communities^3,8^ and, as a result, microbial eco-evolutionary forces are often studied through operationally defined units rather than those on which they act.

In practice, eco-evolutionary dynamics are most studied at the species level. While microbial species are understood conceptually as gene-flow units, practical delimitation has instead heavily relied on the resolution of available methods and technologies, from morphology to 16S rRNA gene similarity and whole-genome comparisons, and been tied to binomial nomenclature-based classification schemes^9–11^. Fostered by the exponential growth of sequencing data, the genomic species concept, which relies on a 95% average nucleotide identity (ANI) threshold^12,13^, has emerged as the standard species-level unit of prokaryotic taxonomy. Yet, this species definition encompasses substantial genetic diversity, including single-nucleotide or gene-content variation, that bear important functional consequences and can organise in finer-grained gene-flow units ^6,14–16^. Whether genomic species or these finer-scale groups within them are biologically meaningful units, i.e. the level at which eco-evolutionary forces act, thus remains an open question.

Addressing this question requires resolving genomic diversity below the species level, without presupposing where the boundaries lie. This is the premise of the reverse ecology framework, which delineates robust eco-evolutionary units by first organizing within-species diversity based on genomic signatures and identifying the ecological drivers of the evolutionary structure afterwards^6–8,17^. Early applications of this framework to isolate genomes demonstrated that strains of *Vibrio* spp. with closely related genomes partitioned into populations with distinct microhabitat preferences^17,18^, while populations of *Ruminococcus gnavus* show differential association with health and inflammatory bowel disease^6^. More recently, genome-wide selective sweeps have been identified as a pervasive mechanism generating ecologically differentiated units in the human gut, which are associated with disease phenotypes^19^. Therefore, these units are not only a finer grained taxonomic resolution of within-species diversity but the scale at which ecology and evolution meet^5^. For semantic consistency, we consider these lineages as subspecies^20^.

Yet, extending those insights to the uncultivated microbial majority^21^ requires moving from cultivation-dependent to cultivation-independent approaches. These strategies can rely on genome-based comparisons^22,23^, marker-gene approaches^24^, or single-nucleotide variant (SNV) profiles^25–31^. In the ocean specifically, single-cell genomics revealed that *Prochlorococcus* harbours distinct sub-lineages adapted to different light conditions^32^. Metagenomic SNV profiling has since uncovered adaptive differentiation in *Pelagibacter* along temperature gradients^33^ as well as population structure in uncultivated species along Baltic Sea salinity gradients^30^. While these studies collectively demonstrate the ecological relevance of subspecies-level resolution in the marine environment, they remain limited in taxonomic scope and geographic scale. As a result, the prevalence of subspecies-level units in the global ocean microbiome is unknown despite its role in global biogeochemical cycles and food webs^34^.

Here, we leverage thousands of genomes from the Ocean Microbiomics Database^35^ alongside 1,669 globally distributed metagenomic samples to apply the reverse ecology framework at the scale of the global ocean. We identify genomically discrete subspecies in 270 of 413 species spanning 15 phyla, including uncultivated lineages without reference isolate genomes. We characterise how they diverge in a subspecies-specific manner, through gene-specific selection on the core genome and restructuring of variable gene content, and further demonstrate that these subspecies represent ecologically cohesive units through global correlations and case studies. Together, these results suggest that subspecies are prevalent biologically meaningful units of the global ocean microbiome.

## RESULTS

### Discovery of genomically discrete subspecies

Within-species genetic variation arises from mutation and gene flow, and is shaped by drift and selection. Depending on the balance of these forces, a species may retain a single, freely recombining gene pool or split into cohesive, genetically distinct subspecies-level groups^20,36^. The extent to which such discrete groups exist in natural microbial populations, and at what level of genomic divergence they emerge, remains largely unknown. To address this across the global ocean microbiome, we mapped single-nucleotide and gene content variation at scale.

Large-scale analysis of within-species diversity in the ocean microbiome requires reference genomes for species that remain largely uncultivated. To that end, we leveraged a collection of ∼35,000 marine microbial genomes spanning >8,000 species^35^ and filtered for those represented by at least 10 genomes to adequately capture their gene content. For each species, we selected a representative genome based on integrated quality metrics, used all available genomes to build a gene catalogue at 95% nucleotide identity, and determine the species -level core genes (Methods). We mapped within-species variation across 1,669 metagenomes, collected at 295 globally distributed sites, restricting the analysis to species with sufficient coverage (Methods). This yielded 27,111 SNV frequency as well as gene content profiles across 413 species. To test whether SNV profiles vary continuously across samples or group into discrete subspecies, we embedded variant frequencies in a low-dimensional space and used density-based clustering to resolve subspecies structures, independently of *a priori* thresholds or number of clusters (Figure 1). Relative to previously described subspecies discovery pipelines^26,28^, our approach introduces improved variant-calling accuracy, captures non-linear structure, identifies genomically discrete subspecies independently of a threshold, and scales across the global ocean microbiome (Methods).

**Figure 1.**
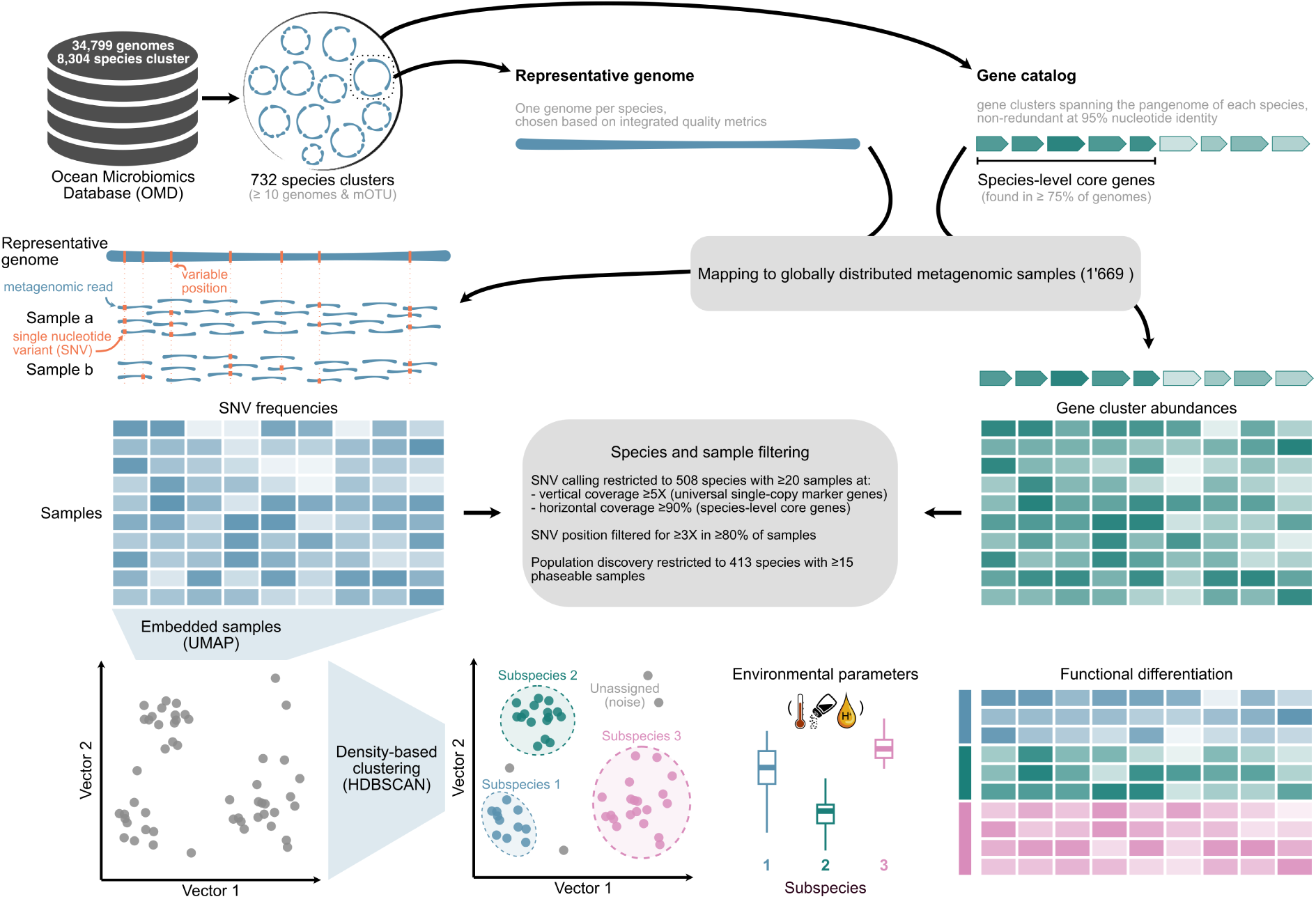
Workflow to infer subspecies from within-species nucleotide variation profiles. Species-level clusters from the OMD were selected if they included at least ten genomes and a mOTU assignment. A representative genome was selected for each species based on integrated quality metrics (Methods) and all genes from all genomes were used to build a gene catalogue and identify species-level core genes (Methods). 1,669 metagenomes from the global ocean were mapped to all representative genomes for SNV profiling as well as gene catalogues for functional profiling. Species were filtered based on the number of samples (at least 20) meeting coverage criteria (Methods) prior to SNV calling. In addition SNVs were retained only if the position was covered at 3X in 80% of the samples. Only species with at least 15 phaseable samples were retained for further analyses. SNV profiles were embedded in two-dimension spaces (UMAP) and clustered with a density-based algorithm (HDBSCAN), which assigns each sample to a subspecies cluster together with an assignment probability. Subspecies can then be tested for associations with environmental parameters (temperature, salinity, oxygen, iron, nutrients, depth) as well as functional differentiation.

We identified 724 genomically discrete subspecies in 270 out of 413 species (Figure 2a). Within-subspecies distances were consistently smaller than between-subspecies distances across a range of independent metrics (Figure 2c-e), including original SNV frequencies, functional content (both gene-cluster and Pfam families), and core gene phylogenies (Methods). Using species core genes alignments, we further found the average nucleotide identity was higher within (mean 99.21%) than between (mean 98.76%) subspecies, although no clear threshold emerged across species (Supp. Fig. S1). Species with subspecies spanned two archaeal and 13 bacterial phyla and accounted for 20.5±9.7% of the relative abundance of marine microbial communities (Supp. Fig. S2). Species without predicted subspecies were detected in fewer samples (Supp. Fig. S2, Wilcoxon test), suggesting that their apparent lack of structures may reflect undersampling rather than a different organisation of within-species diversity. Overall, we found genomically discrete subspecies (median = 2 per species) to be widespread and ecologically important in the global ocean microbiome.

**Figure 2.**
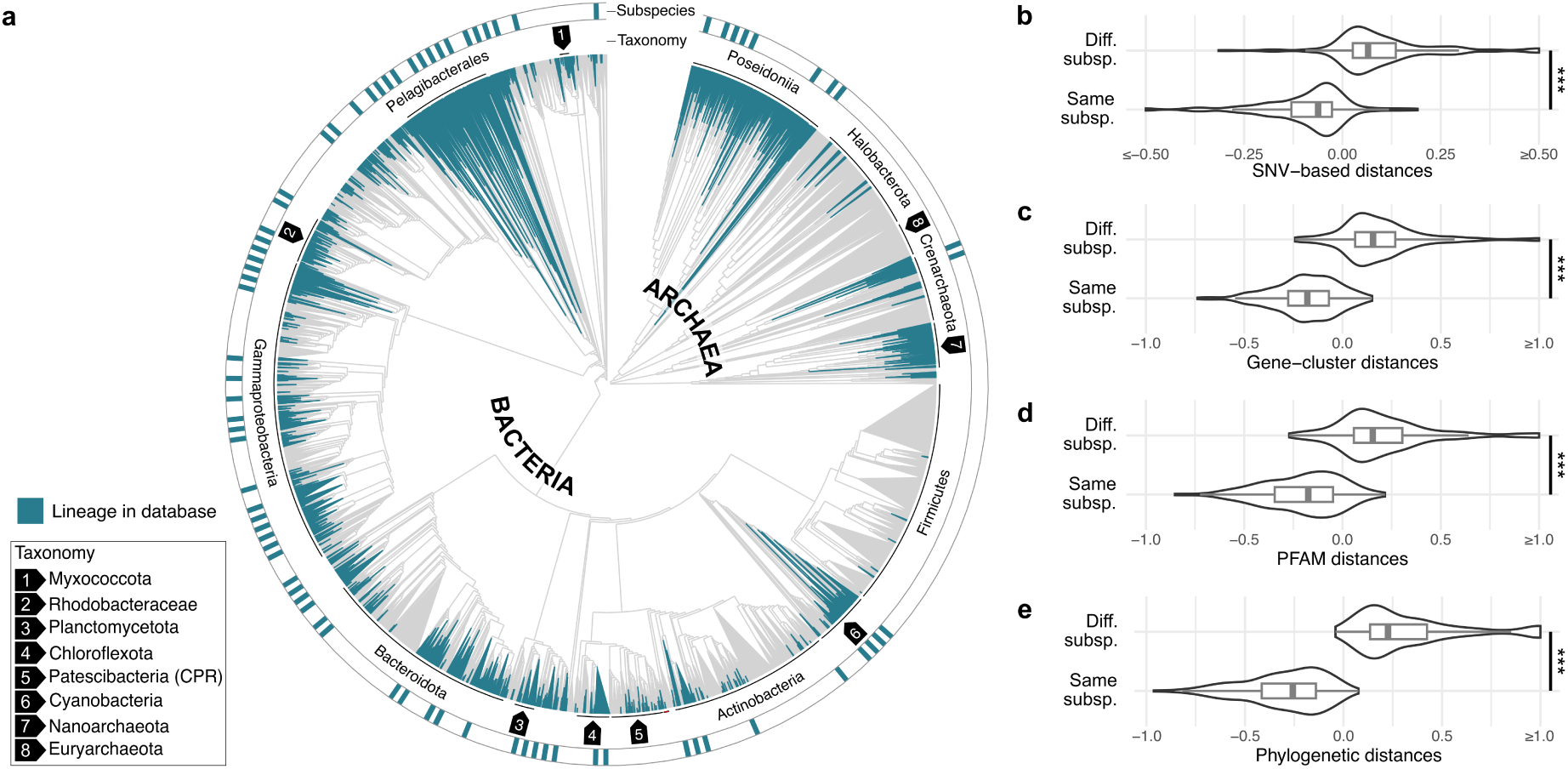
Genomically discrete subspecies are widespread in the ocean microbiome. **a)** Phylogenomic tree of bacterial and archaeal species from the OMD (teal) placed in the GTDB backbone tree (grey). The outer ring marks the species in which genomically discrete subspecies were identified, spanning two archaeal and 13 bacterial phyla; the inner ring delineates major taxonomic groups, labelled around the tree, with less prominent groups numbered and listed in the inset. **b–e)** Distances between samples assigned to the same subspecies (Same subsp.) versus to different subspecies (Diff. subsp.) of the same species, computed on four metrics: **b)** SNV frequency profiles in the original (non-embedded) space, **c)** gene-cluster content, **d)** Pfam family content, and **e)** core-gene phylogenies (Methods). Distances are centred per species and truncated at the axis limits for display. In all four metrics, within-subspecies distances were significantly smaller than between-subspecies distances ([Wilcoxon test]; ***, P < 0.001). Boxes show the median and interquartile range, whiskers extend to 1.5× the IQR, and violins show the full distribution.

### Mechanisms of diversification

Having identified discrete subspecies, we next sought to characterise the mechanisms underlying their divergence, considering two complementary axes: nucleotide-level variation, and restructuring of gene content through gene gain or loss. We investigated each of these axes in turn, before testing the extent to which they are coupled across species.

Signatures of differential selective pressure manifest through both diversifying selection and variation in purifying constraint, and are consistently structured at the subspecies level and concentrated within only a fraction of the core genomic backbone. By analysing the ratio of non-synonymous to synonymous polymorphisms (pN/pS), we found that species’ core genes were, on average, subject to strong purifying selection (Figure 3a), in line with previous observations on marine microbes^37,38^. However, the extent of purifying selection and the distribution of pN/pS values varied across species (Kruskal-Wallis *p* < 2.2×10^−16^) and, in the majority of species tested (n = 200), across subspecies too (adjusted *p* < 0.05). Using a generalised linear mixed-effects model, we found that species (10.5%) and subspecies (1.2%) identity explained only a minor fraction of the variance attributable to the modelled random effects, which was instead overwhelmingly driven by gene identity (88.2%), indicating that selective pressure is primarily a gene-specific property. Consistent with this, only a minor fraction of core genes (0 - 3.7%, n = 4,317) showed evidence of diversifying selection (pN/pS > 1) (Figure 3b), and the vast majority of these signals (90.6%) were either shared by only some subspecies or were subspecies-specific. Similarly, 4.1% (n = 24,093) core genes exhibited a significant subspecies -specific shift in the magnitude of their pN/pS (see Methods). These genes under differential selective pressure were found to span diverse roles, with no convergence on particular KEGG pathways or metabolisms in permutation-based enrichment tests.

**Figure 3.**
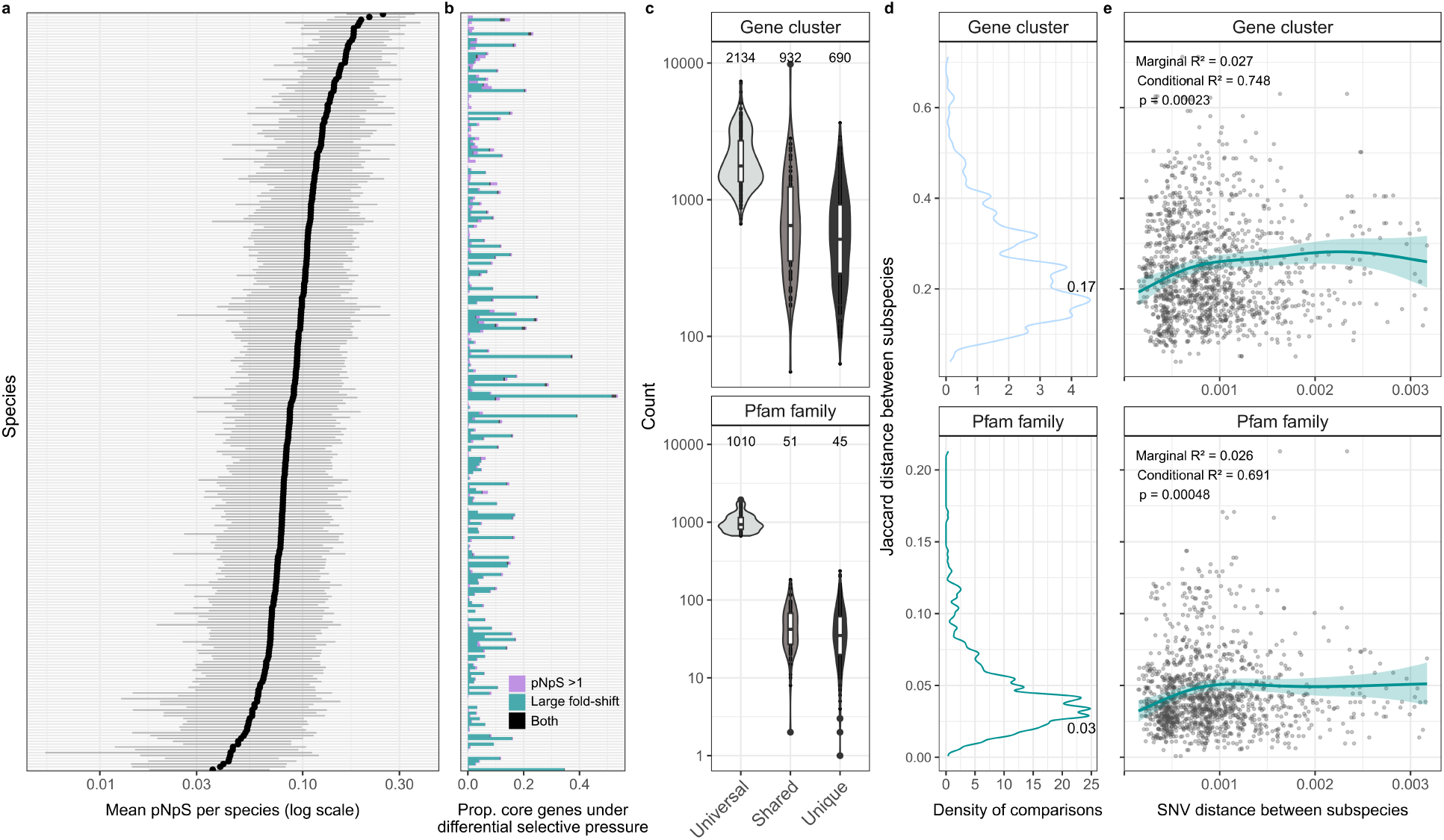
Nucleotide to protein functional repertoire divergence of subspecies. The core genomic content of gene clusters and Pfam families was determined at the subspecies-level after inferring an empirically derived threshold of 99% prevalence across samples. Species core genes were then defined as those shared by all subspecies. **a)** Median (points) and Inter-Quartile Range (IQR, lines) of species core gene clusters across samples. **b)** The proportion of the species core gene clusters either exhibiting pN/pS >1, a significant fold-change in pN/pS between subspecies, or both. **c)** The number of subspecies-core gene clusters and Pfam families that are shared across all (Universal), some (Shared) or specific to a single (Unique) subspecies within a species. Boxplots illustrate the interquartile range along with the median, while the whiskers depict the 1xSD of the IQR. The text labels indicate the mean values. **d)** Density plot illustrating the distribution of Jaccard distances between the core gene cluster and Pfam content of subspecies within the same species. **e)** The Jaccard distances of subspecies’ core genomic content were modelled as a function of the subspecies SNV distances using a generalised additive mixed model (GAM) with species as a fixed effect. Importantly, only the 99th percentile of distance values were modelled, as the sparsity of points at the highest end of the spectrum introduced considerable noise (see Supp. Fig. S4). Points show individual subspecies pairs. The lines and shading show the fitted GAM smooth curve and 95% confidence interval.

Gene-content divergence between subspecies follows a similar pattern, with each subspecies encoding distinct, unique content on top of a shared core backbone. To quantify this, we determined the core genetic content (gene cluster and Pfam families) of each subspecies using an empirically determined cut-off based on prevalence (99%) across samples (Supp. Fig. S3). This revealed that the core genome of subspecies comprised 2705±59 gene clusters, corresponding to 1037±11 distinct Pfam families. The majority of the core genetic content, 70±1.0% of gene clusters and 94±0.2% of Pfam families, was conserved at the species -level. However, on top of this core backbone, each subspecies additionally encoded unique content, comprising 690±34 gene clusters and 45±2 Pfam families (Figure 3c). These differences were further evident through pairwise Jaccard distances between subspecies core genomes, which were more tightly constrained for Pfam families (peak at 0.03) than for gene clusters (peak at 0.17) (Figure 3d), but with large variations across species.

To determine the extent to which these two axes of divergence are coupled, we compared the between-subspecies SNV distances with the Jaccard distances of core genetic content. Using generalised additive mixed models (see Methods), we found that both gene cluster and Pfam family content increased non-linearly with SNV distance before plateauing at higher levels (gene cluster: edf = 3.16, p < 2.2×10⁻¹⁶; Pfam family: edf = 3.24, p < 2.2×10⁻¹⁶) (Figure 3e). However, SNV distance alone explained only a modest proportion of the variance in functional divergence (marginal R² = 0.027 and 0.026 for gene cluster and Pfam family, respectively), while the majority of variation was captured through species identity (conditional R² = 0.748 and 0.691). The weak overall coupling between these two axes likely reflects underlying differences in how gene flux and mutation arise across species. For instance, gene-content turnover is shaped by the mode and rate of horizontal gene transfer and by demographic factors, such as effective population size^39,40^, while mutation rates can vary by orders of magnitude across species^2,41^. Together, these observations indicate that within-species diversification proceeds through both subspecies-specific selection on the core genomic backbone and variable restructuring of gene content, but the coupling between these axes is species-specific.

### Drivers of within-species diversity

Having established that microbial species in the global ocean exhibit genomically discrete subspecies with functional divergence, we sought to explore the drivers of these structures, including gene-flow, selection, and drift^42^. While distinguishing these processes from observational data is challenging, we tested subspecies for associations with environmental parameters and Lagrangian models of oceanic connectivity, which serve as proxies for selection and gene-flow, respectively. An association with environmental parameters implies that subspecies are differentially favoured across conditions, whereas an association with larger connectivity times may reflect geographic barriers to gene flow (dispersal limitation), assuming environmental covariates are accounted for. Under this framing, subspecies showing neither signal may result from unmeasured selection or reflect drift.

We then built species-level multinomial models testing subspecies for association with temperature, salinity, oxygen, iron, nutrients (phosphate, nitrate), connectivity, and depth. Given sparse and heterogeneous *in situ* measurements, we additionally included harmonised environmental parameters from ocean models (Methods) and derived oceanic connectivity from Lagrangian surface transport models^43^. As these models are restricted to surface waters and have variable accuracy across environmental parameters (Methods), we fitted three complementary model types: *in situ* measurements with connectivity in surface waters, modelled parameters with connectivity in surface waters, and depth across the full water column.

We were able to fit environmental models for a total of 142 species. We found significant associations for 88 of these species, with 76 linked to at least one environmental parameter (Figure 4a). In the harmonised models (n = 134), the most frequent associations were found with temperature (a third of the species) followed by connectivity (30.6%), while salinity, nutrients, and iron were each associated with 15–20% of species (Figure 4b,d). Temperature and oxygen are strongly collinear in the surface ocean and could not be disentangled with the data available. Separately, we tested associations with depth for close to 250 species and found significant associations in 24% of cases (Figure 4d,e). However, associations with depth and/or connectivity were often found in species with environmentally structured subspecies. This suggests that selection and dispersal limitation may simultaneously contribute to within-species structures, although disentangling their relative contributions is non-trivial. In addition, connectivity and depth may also capture environmental covariates absent from the model and thus represent unmeasured selection. Conversely, we detected no association for a third of the species. While drift may be one of the factors at play, we tested only a subset of relevant environmental parameters, and additional metadata would likely reduce this fraction. Despite a probable underestimation of environmental associations, subspecies structure in the global ocean microbiome is linked to environmental conditions for half of tested species, with temperature the most widespread among them. Together with adaptive signatures in core genes (pN/pS) and gene-content variation between subspecies, these results suggest that environmental selection is a key driver of genomically cohesive subspecies.

**Figure 4.**
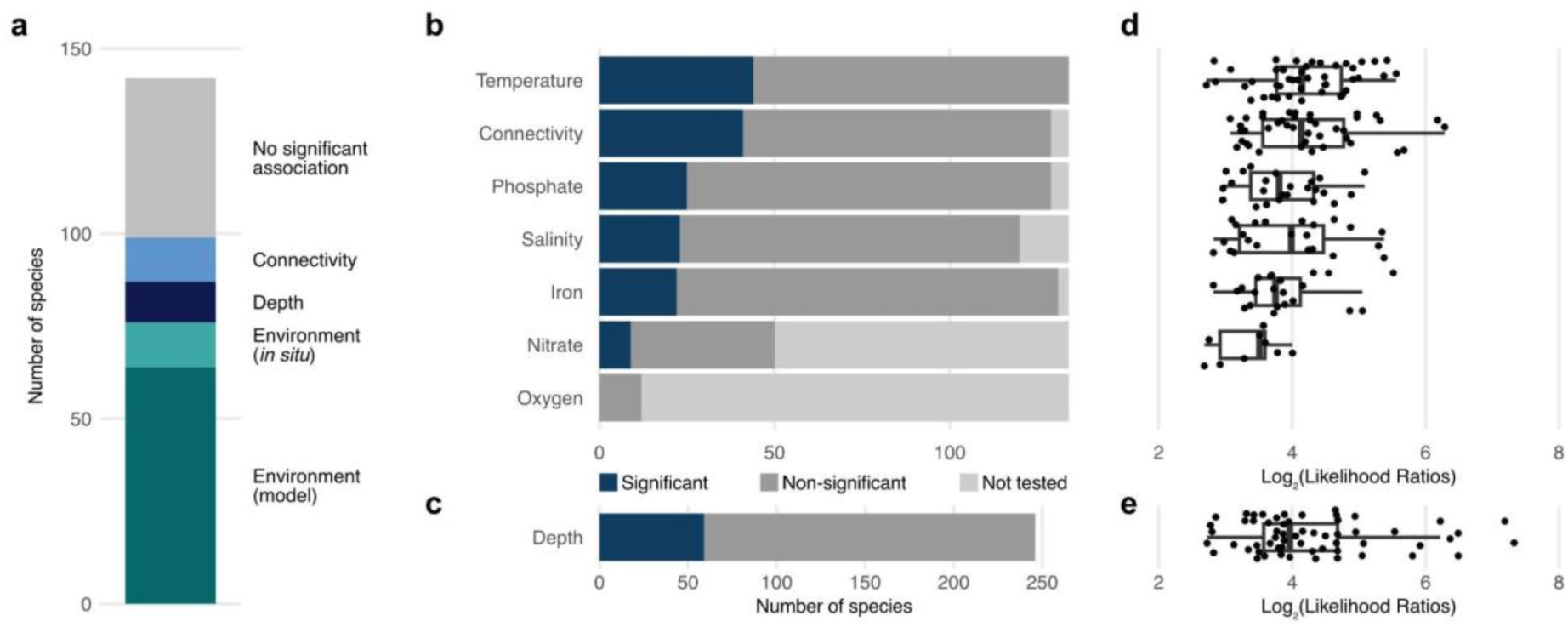
Environmental drivers of subspecies diversification. Subspecies were tested for association with environmental parameters, ocean surface connectivity, and depth using species-level multinomial (Methods). **a)** Summary of the outcome across all fitted models (n = 142 species). Subspecies were associated with at least one environmental parameter in half of the species, using either modelled (dark teal) or in situ (light teal) measurements; with depth but not the environment in 8% of species (dark blue); and with connectivity but neither the environment nor depth in 7% (light blue). No significant association was detected for the remaining third (grey). **b)** Number of species for which each parameter was significantly associated with subspecies structure (dark blue), not significantly associated (dark grey), or not tested (light grey) owing to collinearity with other variables (Methods) in the multinomial models with harmonised environmental parameters and connectivity in surface water (n = 134 species). Temperature was the most frequent association (a third of species), followed by connectivity (just under 30%), with phosphate, salinity, and iron each associated with 15–20% of tested species. Oxygen could not be disentangled from temperature owing to their strong collinearity in the surface ocean and was therefore not tested. **c)** As in b, for the depth models fitted across the full water column (n = 250), with significant associations in 24% of species. **d–e)** Effect sizes of the significant associations, expressed as log₂ likelihood ratios between the full and reduced models for the corresponding parameters. Each point is one species; boxes show the median and interquartile range, and whiskers extend to 1.5× the IQR.

### Subspecies as eco-evolutionary units

To illustrate the coupling between environmental conditions and subspecies genomic divergence, we present two contrasting case studies: one structured by spatial and depth-related gradients, and another by local variation in iron availability. For each, we integrate signals across nucleotide, codon, and gene-content levels to characterise the underlying mechanisms of diversification.

### Subspecies within a SAR324 species are structured over spatial gradients

As a first example, we focused on Arctic96AD-7 sp002082305, an uncharacterised species within the Arctic96AD-7 genus (SAR324 Clade), that was distributed across all major ocean basins and latitudinal zones. Across this broad distribution, we identified three subspecies (Figure 5a) with distinct biogeographical structuring, partitioned primarily by depth (multinomial regression LogOdds ratio = 160.53) and latitude (Figure 5b): Subspecies_0 (deep waters, temperate/tropical), Subspecies_1 (epipelagic, polar), and Subspecies_2 (epipelagic, tropical/temperate). We therefore asked whether this spatial partitioning is supported by adaptive changes at the genomic and functional level.

**Figure 5.**
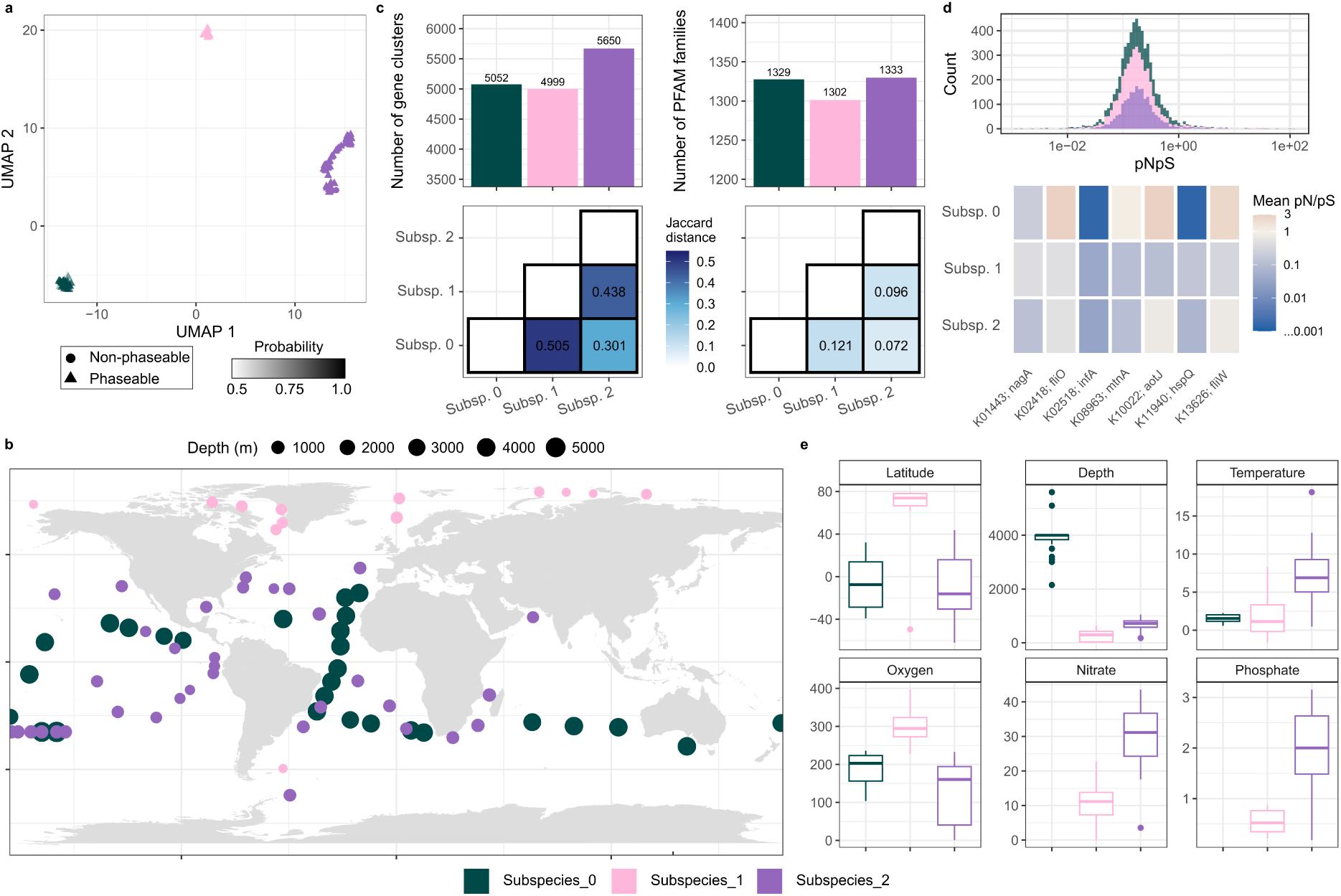
Subspecies within a species of Arctic96AD-7 occupy discrete niches across latitudinal and depth gradients. **a)** UMAP embedding of SNV distances between metagenome samples, which were grouped into three distinct subspecies through HDBSCAN (Methods). The transparency of the points represents the probability of assignment to a given subspecies, while the shape determines whether it was used for subspecies discovery (phaseable) or assigned a posteriori (non-phaseable). **b)** The biogeographical distribution of subspecies, with points representing individual metagenome samples, and their size reflecting the depth below the surface. **c)** Barplots represent the number of gene clusters and Pfam families identified within the core genome of each subspecies - based on an empirically determined threshold of 99% prevalence across samples. The heatmap illustrates the Jaccard distance between the core genome of each subspecies. **d)** The histogram represents the distribution of pN/pS values across the core gene clusters of each subspecies (with each gene cluster value derived from a mean across samples). pN/pS values were subsequently scaled to the KEGG ortholog level based on the mean of the respective gene clusters. A subset of KEGG orthologs that either exhibited pN/pS >1 or the greatest magnitude in difference between subspecies were selected for inclusion in the heatmap. **e)** The distribution of geographical or physicochemical parameters across metagenomes phased to each subspecies - only measurements collected *in situ* with the original sample are included.

The subspecies exhibited clear, consistent signatures of genomic divergence. The discrete subspecies structure identified through SNV compositions was corroborated by phylogenetic analysis of the species’ core genome (Supp. Fig. S5), codon usage patterns within ribosomal proteins (Supp. Fig. S5), and gene-content composition (Figure 5c). Across each of these, Subspecies_0 and Subspecies_2 were also consistently more similar to one another than either was to Subspecies_1, indicating the latter as the most divergent lineage. This was further evident on a protein function level, with Subspecies_1 encoding a greater functional repertoire, comprising >60 additional Pfam families (n = 1,387) (Figure 5c).

These genomic differences encompassed signatures of adaptation to different environmental contexts. Overall, the species was found to be metabolically versatile, encoding both the core machinery for carbon fixation via the CBB cycle and a broad capacity for heterotrophic growth on organic compounds as carbon and sulfur sources, in line with a mixotrophic or facultative chemolithoauthotrophic lifestyle previously proposed for SAR324. Against this shared backbone, which was under purifying selection (Figure 5d), each subspecies carried distinguishing metabolic and nucleotide selective signatures. For Subspecies_0, these included a cyanophycin synthesis/degradation system (*cphA*/*cphB*), mercuric ion transporter (*merC*) and urea-proton symporter (DUR3), alongside elevated pN/pS on arginine/ornithine transport (*aotJ*) and methionine salvage (*mtnA*) and stronger purifying selection on genes involved in stress response (*hspQ*) (Figure 5d), in line with adaptation to the reduced, heterogeneous nutrient availability of the deep ocean. In contrast, Subspecies_1 harboured an expanded repertoire of carbohydrate degradation machinery, including for lyxose (*lyxA*/*lyxD*) and xylose (*xylA*), as well as a sensory rhodopsin (*sop*) and an ECF sigma factor (*rpoE*), along with reduced purifying selection in N-acetylglucosamine catabolism (*nagA*), reflecting adaptation to the variable organic matter pool of the Arctic Ocean. Meanwhile, Subspecies_2 additionally encoded a near-complete dissimilatory sulfate reduction pathway (*dsrAB, dsrMKOP*), building on the upstream adenylylsulfate reductase (*aprAB*) shared with Subspecies_0.

Together, these results illustrate that within-species diversification in Arctic96AD-7 sp002082305 has culminated in the emergence of three subspecies with distinct nutrient-acquisition and stress-response strategies, adapted to discrete environmental niches (Figure 5e) across vertical and latitudinal gradients in the ocean.

### Subspecies within a Gammaproteobacteria species are structured by iron availability

As a second example, we examined AAA300-D14 sp002715065, an uncharacterised species within the AAA300-D14 genus (*Porticoccaceae; Gammaproteobacteria*), which comprised three subspecies (Figure 6a) distributed across tropical regions of the Atlantic, Pacific and Indian Oceans (Figure 6b). Despite this more restricted distribution, *in situ* measured iron concentrations were a significant predictor of subspecies occurrence (multinomial regression, LogOdds Ratio = 13.6). Therefore, we investigated whether the subspecies exhibit distinct genomic and functional signatures consistent with adaptation to contrasting iron regimes.

**Figure 6.**
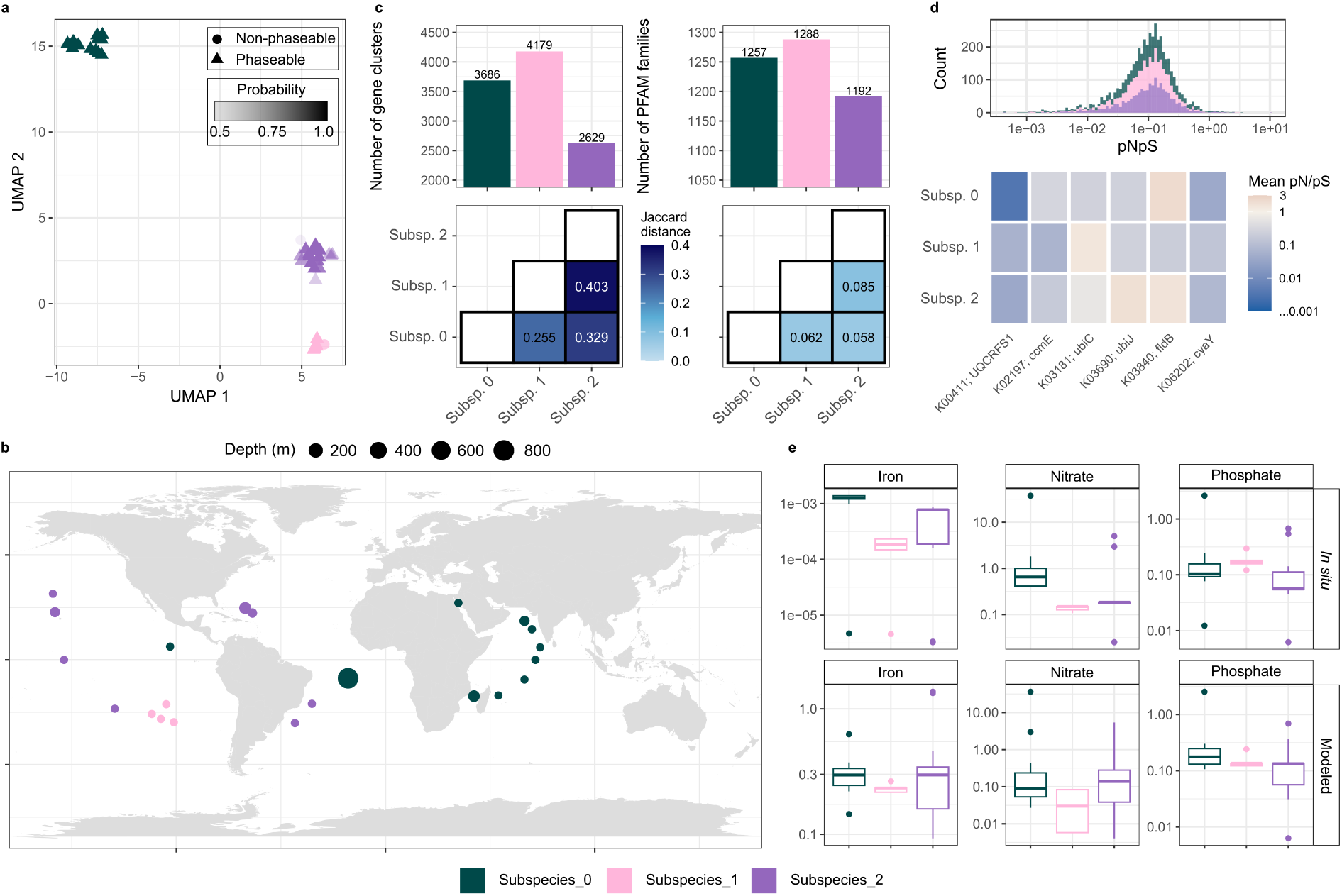
Subspecies within a species of AAA300-D14 genera occupy discrete niches in relation to iron availability. **a)** UMAP embedding of SNV distances between metagenome samples, which were grouped into three distinct subspecies through HDBSCAN (Methods). The transparency of the points represents the probability of assignment to a given subspecies, while the shape determines whether it was used for subspecies discovery (phaseable) or assigned a posteriori (non-phaseable). **b)** The biogeographical distribution of subspecies, with points representing individual metagenome samples, and their size reflecting the depth below the surface. **c)** Barplots represent the number of gene clusters and Pfam families identified within the core genome of each subspecies - based on an empirically determined threshold of 99% prevalence across samples. The heatmap illustrates the Jaccard distance between the core genome of each subspecies. **d)** The histogram represents the distribution of pN/pS values across the core gene clusters of each subspecies (with each gene cluster value derived from a mean across samples). pN/pS values were subsequently scaled to the KEGG ortholog level based on the mean of the respective gene clusters. A subset of KEGG orthologs that either exhibited pN/pS >1 or the greatest magnitude in difference between subspecies were selected for inclusion in the heatmap. **e)** The distribution of physicochemical parameters across metagenomes phased to each subspecies, including measurements derived *in situ* and from models (Methods).

Across a shared metabolic backbone of chemoheterotrophy, subspecies exhibited notable signatures of genomic divergence. Notably, Subspecies_0 and Subspecies_1 harboured greater protein functional repertoires (n = 1,257 and 1,288, respectively) and were more similar in their genetic content (Jaccard distances: Gene cluster = 0.255 and Pfam families = 0.062), while Subspecies_0 was more streamlined and distinct (Figure 6c). Amongst the additional protein functions observed in Subspecies_1, we identified a number of distinctions associated with iron-acquisition machinery, including a uniquely encoded heme substrate-binding protein and permease (*hmuT*, *hmuU*), an iron-complex ABC transporter ATP-binding domain (*ABC.FEV.A*) and a Fe(3+)-dicitrate transporter (*fecA*), alongside a higher copy number of the TonB-associated *exbD* component. This targeted expansion of iron-acquisition machinery in Subspecies_1, which occupies comparatively iron-limited conditions in the South Pacific Gyre, highlights gene-content restructuring as one route of adaptation to distinct iron regimes within this species.

Signatures of nucleotide-level selection further reinforced these patterns, with a subset of genes exhibiting clear subspecies-specific selective pressure centred on iron-dependent metabolism, despite no significant subspecies-level structure across the core genomic backbone (Figure 6d). Flavodoxin II (*fldB*), which substitutes for the iron-requiring ferredoxin under iron limitation, was under strong purifying selection in Subspecies_1 (pN/pS = 0.21) but positive selection in Subspecies_0 and Subspecies_2 (pN/pS = 2.15 and 1.84), directly reflecting its contrasting essentiality across the iron gradient. Similarly, genes involved in iron-sulfur cluster (*cyaY*) and heme cofactor biogenesis (*ccmE*) also show subspecies-specific differences in purifying selection, particularly across Subspecies_0 and Subspecies_1 (Figure 6d), pointing to broader selective pressure on iron-related cofactor biosynthesis.

Together, these results show that subspecies within AAA300-D14 sp002715065 are structured along a fine-scale gradient of iron availability (Figure 6e), aligned with distinct, targeted signatures of genomic divergence. This concordance between environmental regimes and specific, adaptive genomic changes illustrates how ecological differentiation can emerge through specific acquisition and regulatory functions within a shared heterotrophic lifestyle.

## DISCUSSION

Much of the eco-evolutionary dynamics shaping microbial life is thought to unfold below the species level, yet this scale of diversity has remained difficult to resolve across the breadth and complexity of natural communities. Here, we resolve within-species diversity in 270 microbial species across the global oceans, using a scalable, cultivation -independent framework that infers subspecies structure empirically from metagenomic SNV profiles. We demonstrate that these subspecies represent genomically discrete and ecologically cohesive units whose divergence, in many cases, is associated with distinct environmental conditions. Thus, these findings support subspecies as a fundamental level of biological organisation at which ecological and evolutionary processes jointly shape microbial diversity.

### A cultivation-independent, scalable framework for subspecies-level inference

Our framework builds on, but addresses key constraints in previous approaches for profiling within-species diversity. While recent developments have overcome the dependence on cultivated isolates and extended population-level analyses to the uncultured microbial majority^30^, resolving population structure across thousands of samples and hundreds of species requires an analytical approach that is both scalable and able to accommodate variation in the extent of within-species divergence. We addressed this by combining metagenomic SNV profiling with low-dimensional embedding and density-based clustering, which enables subspecies to be inferred without imposing pre-determined divergence thresholds or assumptions about diversification rates across species^27,44^. As this process captures dominant compositions rather than exact clonal identities, it resolves diversity at a coarser-grained resolution than strains, and at a level more comparable to MAGs. As such, it complements, rather than replaces, strain-resolution tools, such as InStrain^27^, ZipStrain^45^, or TRACS^46^ or frameworks centred on selective sweep detection^7,19^. Furthermore, by conducting this analysis on a per-species basis, our framework remains computationally tractable at scale, and allows new samples to be projected directly into existing embeddings and classified against previously delineated subspecies. Together, these properties allow subspecies structure to be inferred empirically and at scale across the full breadth of microbial diversity captured in metagenomic data.

While our framework offers several key advances, it retains some dependencies that impose practical constraints. Namely, it still requires a genomic representative of sufficient quality, i.e. high completeness, for each species. In addition, as subspecies structure is inferred from metagenomic read alignment, the sensitivity of the approach is tied to the number of samples available and the coverage of a given species within them. Species that are consistently rare or present at low abundance are thus unlikely to yield sufficient signal for subspecies discrimination. Nevertheless, given the rapid and ongoing expansion of environmental sequencing efforts and the accelerating recovery of high-quality MAGs^47,48^, the fraction of species for which this framework is applicable will continue to grow in the future.

### A spectrum of within-species diversification in the global ocean microbiome

Consistent with the view that within-species diversification is a dynamic and continual process rather than a series of discrete evolutionary transitions^5,36,49,50^, our SNV-based framework recovers subspecies spanning a broad spectrum of divergence, from subtle nucleotide -level variation to substantial restructuring of gene content and function.

At the nucleotide level, our analysis of pN/pS patterns points to a mode of divergence consistent with early-stage differentiation, in which localised adaptation to distinct ecological contexts leaves detectable signatures within the otherwise shared genomic backbone of a species. Signatures consistent with positive selection were restricted to only a small fraction of the core genome and were predominantly subspecies-specific. Such localised signals are reflective of gene-specific selection that occurs without disrupting the shared genomic background or ongoing gene flow between subspecies^17,23,51,52^. While elevated pN/pS is also compatible with relaxed purifying or balancing selection^53^, its restriction to a small, subspecies-specific set of genes argues against genome-wide relaxation, and instead points toward selection operating locally and unevenly across the genome^17^. In further support of this, the functional roles that such genes were associated with spanned diverse cellular processes, and lacked convergence across subspecies and species. Together, these observations indicate that this mode of divergence primarily acts in a gene-specific manner within individual subspecies, and is likely underpinned by variation in selective pressure across different environmental contexts.

A second mode of divergence involves shifts in gene content that, in some cases, manifest in broader reconfiguration of metabolic capacities. This mode of divergence varied considerably in magnitude across species, but was nonetheless a pervasive feature that, like nucleotide-level selection, was consistently structured at the subspecies level. The marked variability observed likely reflects how gene flow occurs and is shaped across species, from the mode and rate of recombination and horizontal gene transfer to demographic factors, such as effective population size, each varying according to differences in lifestyle and subspecies history^39,40,54^. As subspecies adapt to new ecological contexts, the selective value of specific functions can shift, favouring the retention of newly acquired genes, or the loss of others, in a manner that can directly underpin adaptation at the within-species level^1,5,7^. Such events have been previously exemplified for nutrient acquisition genes among *Prochlorococcus* ecotypes^55,56^ and phage-defence elements among wild *Vibrio* populations^57^, and were evident across the subspecies identified in this study, with the case studies providing clear examples tied to distinct nutrient and trace -metal regimes. Gene-content restructuring thus represents a distinct, ecologically consequential route of subspecies divergence only partly coupled to nucleotide-level change.

These two axes of divergence exhibit marked differences in their coupling across species. This pattern likely reflects their complementary roles within a single diversification process, with gene content restructuring plausibly acting as the trigger of ecological differentiation and the consequent restriction of gene flow, and genome-wide nucleotide divergence accumulating more gradually and downstream, tracking the strength and duration of this isolation. Under this interpretation, the variable coupling observed would reflect differences in the time since subspecies divergence, with the pace and mode of diversification itself differentially shaped by ecology, connectivity, and historical contingency across species^5,50^. Over evolutionary timescales, the progressive accumulation of divergence through these routes will give rise to recombination barriers that further restrict gene flow, potentially seeding the emergence of distinct species^58–61^. The recovery of subspecies that span this within-species spectrum thus offers a window into this continuous process, capturing ecologically driven diversification as it unfolds and shapes long-term evolutionary outcomes.

### Genomically discrete and ecologically cohesive subspecies as biologically meaningful units

The subspecies identified through our framework represent an emergent level of organisation within microbial species. At finer scales, species are composed of transient genotypes that fluctuate dynamically under the effects of dispersal, competition, and frequency-dependent pressures such as phage predation and grazing^15,16,62–65^. In contrast, the recurrent genomic compositions identified by our framework integrate across this finer-scale variation, capturing groups of related individuals that have accumulated sufficient adaptive advantage to repeatedly dominate a particular ecological context. Subspecies may thus encompass local genotype -level turnover while retaining a shared genomic background shaped by a common selective environment. It is this coupling of genomic discreteness and ecological cohesiveness, rather than any fixed degree of sequence divergence, that defines these subspecies as biologically meaningful units^7,49^. They occupy an intermediate position along the continuum between strains and species, representing a snapshot of ongoing diversification wherein ecological selection has generated reproducible genomic structure. The prevalence of such structure across the global ocean microbiome suggests that this is a fundamental level at which ecological and evolutionary processes jointly drive divergence and structure microbial diversity in nature.

## METHODS

### Species selection, representative genomes, gene catalogues, and core genomes

We leveraged the Ocean Microbiomics Database (OMD)^35^, which contains >34,799 metagenome-assembled^66^, single-cell amplified^16^, and isolate^67^ genomes representing >8,300 species-level clusters (95% ANI), most of them uncultivated. To ensure that the gene content of each species was adequately represented, we retained only species clusters comprising at least 10 genomes. For the purpose of downstream analyses, we further filter for species clusters with mOTUs assignment.

For each retained species, we selected a representative genome based on completeness and contamination estimates, contiguity, genome size, and presence of tRNAs and 16S rRNA genes, extending a previous scoring strategy^68^. In particular, we sought to account for differences in, *e.g.*, completeness or genome size between species that would otherwise contribute unevenly to the final score across species if not rescaled. As such, we used the following strategy:

Score = 0.5 × quality index + 0.3 × contiguity index + 0.1 × size index + 0.05 × rRNA index + 0.05 × tRNA index, with:

– quality index = rescale(completeness – 5 × contamination / max(quality index of a species))
– size index = genome size / max(genome size of a species)
– contiguity index = rescale(1 – exp(–5 × N50 / genome size) / max(contiguity index of a species))
– rRNA index = 1 if complete, 0.5 if partial, 0 otherwise
– tRNA index = min(number of tRNAs / 20, 1)

With rescale a function that projects a continuous distribution into [0, 1].

Whereas the representative genome provides the reference against which nucleotide variation is called, it captures only a fraction of the gene content present across a species. To represent the full functional repertoire of each species, we additionally constructed a species-level gene catalogue at 95% nucleotide identity spanning all genes from all genomes of the species by subsetting the OMD gene catalog^35^.

The species core genome was defined from these catalogues as the set of gene clusters present in more than 75% of the genomes of a species when accounting for incompleteness. This genome-derived definition is independent of metagenomic detection and is used for horizontal coverage filtering (below).

### Metagenomic sample processing, mapping, and profiling

We leveraged the previous set of metagenomes associated with the OMD and added 631 metagenomes from the *Tara* Oceans larger size fractions^69^, for a total of 1,669 metagenomes collected at 295 globally distributed sites. Raw reads were processed as in Paoli et al.^35^. Briefly, sequencing reads from all metagenomes were quality filtered using BBMap^70^ (v.38.71) by removing sequencing adapters, removing reads that mapped to quality control sequences (PhiX genome) and discarding low-quality reads using the parameters trimq=14, maq=20, maxns=0 and minlength=45. Downstream analyses used merged, unmerged paired and single quality-controlled reads (bbmerge.sh minoverlap=16). All metagenomic samples were mapped against the >8,000 representative genomes to allow for competitive mapping. Alignment files were filtered to retain only alignments of at least 45 bp at 95% identity using a custom script. The resulting BAM files were used to produce Anvi’o^71^ profile databases (using parameters --min-coverage-for-variability 3 and --profile-SCVs) using the representative genomes as the contig database. Gene catalogue profiles were generated by subsetting the OMD gene catalogue profiles^35,72^ on a species-specific basis.

### Selection of species–sample sets for variant calling

Computing SNV profiles for all species across all samples is computationally prohibitive, and uninformative for species–sample combinations in which the species is absent or too sparsely covered for allele frequencies to be estimated reliably. We therefore restricted variant calling using two complementary coverage criteria selecting only species detected in at least 20 samples with enough vertical and horizontal coverage.

Vertical coverage was assessed from pre-computed mOTUs^35,72^ profiles, which estimate species abundance from universal single-copy marker genes. For each species we identified the matching mOTU and filtered for samples where the species was detected at ≥5× coverage. Samples from which a MAG of that species had been reconstructed were additionally retained even where mOTU coverage fell below this threshold.

Horizontal coverage was assessed against the species core genome as defined above. Based on the gene catalogue profiles, species–sample pairs were retained only if at least 90% of the species core genes were detected, ensuring that downstream inference was not driven by partial genome recovery.

### SNV calling and variability profiles

For each representative genome, we generated a contigs database and merged profile databases across all retained samples. The merged profile database was used to generate variability profiles with the Anvi’o SNV engine (anvi-gen-variability-profile, parameters --engine NT --quince-mode).

Variable positions were filtered according to Van Rossum et al.^28^. First, we retained those covered at least three times in at least 80% of samples, restricting the analysis to positions covered consistently across the sample set. Samples were then partitioned into phaseable and non-phaseable subsets. A sample is considered phaseable for a given species if at least 80% of its variable positions carry an allele at a frequency above 80%, i.e. if the majority of core-genome variable positions are near fixation and the sample can therefore be treated as dominated by a single genotype rather than a mixture. Species with fewer than 15 phaseable samples were excluded from subspecies discovery.

### Dimension reduction and subspecies delineation

Subspecies were delineated from the SNV frequency profiles following the general logic of the metaSNV2 pipeline^28^, with two modifications: samples were embedded using a non-linear dimension reduction (UMAP) rather than a linear projection, and groups were delineated by density-based clustering (HDBSCAN), so that neither a similarity threshold nor an expected number of clusters had to be specified in advance.

For each species, the embedding and clustering models were fitted on phaseable samples only as for metaSNV2^28^, since allele frequencies in non-phaseable samples reflect mixtures of genotypes and would blur the structure being resolved. Non-phaseable samples were subsequently projected into the fitted models, so that they contribute to downstream analyses without influencing the delineation itself.

UMAP^73^ hyperparameters depend on dataset properties including sample number and must be set case by case. To retain throughput across hundreds of species, each embedding was fitted 30 times across the hyperparameter space. For each candidate embedding we computed trustworthiness for k neighbours (the retention of distances from the original space) with k set to 30% of the samples available for that species, and selected the embedding maximising this criterion.

The optimal embedding was then clustered with HDBSCAN^74^, a density-based algorithm that is robust to clusters of unequal size, shape and density and that assigns samples not belonging to any dense region as noise rather than forcing their assignment. The hyperparameter space was again explored over 30 iterations, with the optimal clusterer selected by the internal DBCV criterion.

Non-phaseable samples were projected into the fitted UMAP embedding and assigned to clusters using the optimal HDBSCAN model. Samples not assigned to any cluster were retained as noise and excluded from analyses requiring subspecies membership. Fitted models were saved and can be used to classify new samples.

### Benchmarking subspecies

Because subspecies were delineated in the embedded space, we verified that they were also discrete under independent metrics computed without reference to the embedding. For each species, pairwise distances between samples were computed in (i) the original, non-embedded SNV frequency space (euclidean distances of allele frequencies), (ii) the presence/absence of core gene clusters (Jaccard distances), (iii) the presence/absence of core Pfam families (Jaccard distances), and (iv) core-gene phylogenies (see *Phylogenetic reconstruction of within species environmental diversity*). Sample pairs were categorised as within-subspecies or between-subspecies according to their subspecies assignment. Because baseline divergence differs between species, distances were centered prior to pooling, and differences between within- and between-subspecies distances were assessed using a Wilcoxon rank-sum test, separately for each metric.

### Environmental parameters and connectivity

Subspecies membership was tested for association with temperature, salinity, oxygen, nitrate, phosphate, and iron, alongside oceanic connectivity and with depth independently. Because in situ measurements are sparse and unevenly distributed across the metagenome collection, we additionally compiled harmonised values for these parameters from ocean models (Add Dominic’s reference) matched to each metagenome by station. Modelled and in situ values were compared across all samples for which both were available. As the models describe surface conditions, the agreement between the two sources degrades with depth; we therefore evaluated the correlation between modelled and in situ temperature as a function of an increasing depth cut -off within the epipelagic layer, and restricted all environmental models to samples shallower than 100 m (Supp. Fig. S6).

Connectivity between samples was derived from Lagrangian modelling of ocean surface transport, with minimum connectivity times computed using the Dijkstra algorithm as previously published^43^. The latitude and longitude of each metagenome were mapped onto the tiles used in that model, and the connectivity time between two samples was approximated by the connectivity time between their respective tiles. Samples for which connectivity times were missing for the majority of partners (>1,000 pairs) were excluded, and remaining missing values, as well as connectivity times exceeding 4,000 years, were set to a common ceiling of 8,000 years, such that unreachable and very distant sample pairs are treated as maximally disconnected. The resulting distance matrix was projected by principal coordinate analysis (PCoA; *ape::pcoa*), and the first two axes were retained as sample-level explanatory variables representing connectivity^30^.

### Multinomial modeling

For each species, subspecies membership was modelled as a categorical response using multinomial logistic regression (*nnet::multinom* in R; maximum 10,000 iterations). Samples not assigned to any subspecies by HDBSCAN^74^ (noise points) were excluded, as were samples with missing values for any modelled variable. Subspecies represented by fewer than seven samples were dropped, and a species was modelled only if at least two subspecies remained; the subspecies with the lowest index was set as the reference level. All continuous predictors were standardised (zero mean, unit variance) prior to fitting. Collinearity among predictors was addressed by computing the pairwise correlation matrix within each species and removing one member of any pair with an absolute correlation above 0.75. As temperature and oxygen are strongly collinear in surface waters, oxygen was removed from most species’ models under this criterion, and its associations could therefore not be evaluated independently of temperature.

Three complementary model types were fitted per species. First, an environmental model comprising the six harmonised (modelled) parameters together with the two connectivity axes, restricted to epipelagic samples shallower than 100 m. Second, an equivalent model using in situ rather than modelled measurements, under the same sample restriction. Third, a depth model fitted to depth alone across the full water column, thereby retaining samples excluded from the surface models.

The contribution of each variable was assessed by comparing the full model to a reduced model from which that variable had been dropped, using a likelihood-ratio test (connectivity was dropped as a single term comprising both PCoA axes). For depth, the model was compared against an intercept-only null model. P-values were corrected for multiple testing across species and variables using the Benjamini-Hochberg procedure, and associations were considered significant at an adjusted P ≤ 0.05. Effect sizes are reported as log₂-transformed likelihood-ratio statistics. Variables removed from a species’ model owing to collinearity, or absent from its metadata, were recorded as not tested rather than as non-significant.

### Defining the core genome of each subspecies

To empirically determine the core genome of each subspecies, we used information on the distribution of gene clusters across samples. Specifically, we calculated the prevalence of each gene cluster across each subspecies’ assigned samples. Prevalence values were binned at 1% intervals and the first and second inflection point was identified based on the largest shifts in counts across bins. The first inflection point was consistently observed between the 98-99% prevalence bins, a pattern that also held when assessing Pfam families (Supp. Fig. S3). Thus, the subspecies core genome was defined as gene clusters and Pfam families detected in ≥99% of subspecies’ samples. This sample-based definition is distinct from the genome-based species core genome defined above, which is used for coverage filtering.

### Phylogenetic reconstruction of within species environmental diversity

For each species, we determined a core set of single-copy marker genes defined as the Clusters of Orthologous Groups (COGs) that were identified as a single copy in all species’ samples that passed the coverage filter (≥90% species core genome detected). This yielded a species-specific single-copy core gene set of 555–1,266 genes (median ∼700). Each core gene was aligned across samples with MAFFT v7.505 (parameters: --auto), and the per-gene alignments were subsequently concatenated into a per-species nucleotide supermatrix.

Per-species maximum likelihood phylogenetic trees were reconstructed from the nucleotide supermatrices using IQ-TREE2 v2.2.2.7 under the best-fit substitution model selected by ModelFinder (parameters: -m MFP), with 1000 ultrafast bootstrap replicates (parameters: -B 1000). This yielded trees for 207 species.

The patristic distances (tip-to-tip) were computed between all pairs of metagenome samples within each species tree, and the distances were summarised according to representing a within- or between-subspecies comparison. As baseline divergence differs between species, each distance was normalised by that species’ mean pairwise distance before pooling across species. Within-versus between-subspecies distances were compared using a Wilcoxon rank-sum test.

As an independent, alignment-based insight into subspecies divergence, we calculated the pairwise nucleotide identity between metagenome samples within each species. For each pair of samples, nucleotide identity was defined as the proportion of matching positions over all sites at which both samples carried an unambiguous, non-gap base (pairwise deletion). Pairs were categorised as representing a within- or between-subspecies comparison and, as above, normalised per species and compared using a Wilcoxon rank-sum test.

### Analysis of nucleotide-level selective pressure

To assess differences in nucleotide-level selective pressure, we calculated non-synonymous to synonymous polymorphism ratios (pN/pS) for each gene cluster in each sample. For this, we focused on the gene clusters that were present in the core genome of all subspecies within each species, i.e. conserved at the species level. To determine at what level pN/pS typically varied, we fitted a generalised linear mixed-effects model to pN/pS with a Tweedie error distribution and log link (*glmmTMB* package in R), with random intercepts for species, subspecies nested within species, and gene cluster identity. A Tweedie distribution was selected due to a large number of pN/pS values of 0 along with a continuous distribution of positive values. Differences in the variance attributable to each random-effect term were determined from the corresponding conditional-model variance components. To further explore patterns, we then calculated mean pN/pS values for each gene cluster at the species and subspecies level. Differences in the distribution of mean pN/pS values across species and subspecies within species were assessed using a Kruskal-Wallis rank-sum test, with effect size quantified as epsilon-squared.

To identify potential adaptive signals, we extracted gene clusters that either showed evidence of diversifying selection (mean pN/pS >1 in at least one subspecies) or significant fold-change in pN/pS between subspecies of the same species. For the latter, we only considered gene clusters detected in at least three samples of at least two subspecies, and determined fold-change as the ratio between the highest and lowest mean pN/pS across subspecies. To test whether the fold-change was significant, we used a permutation test in which sample-to-subspecies assignments were randomly shuffled within each gene cluster (500 permutations), recomputing the fold-change under each permutation to build a null distribution. Empirical p-values were calculated as the proportion of permutations in which the null fold-change equalled or exceeded the observed value, followed by multiple-testing correction using the Benjamini-Hochberg algorithm.

The KEGG ortholog annotations of gene clusters showing evidence of diversifying selection, and separately of those showing a significant fold-change, were used to retrieve the corresponding KEGG pathway and metabolism categories with the *KEGGREST* package in R. To test whether either set of genes was enriched for particular pathways or metabolism categories, a species-stratified permutation test was used. For each permutation, gene category labels (none/subspecies-specific/species-wide or significant/non-significant for the fold-change analysis) were randomly shuffled among a species’ own core genes, preserving the number of genes within each category, and the number of genes falling within a given pathway/metabolism under this randomisation was recorded across 500 permutations to build a null distribution. Empirical p-values were calculated as the proportion of permutations in which the null count equaled or exceeded the observed count for each pathway/category, followed by multiple-testing correction using the Benjamini-Hochberg algorithm.

### Assessing gene-content and functional divergence across subspecies and its coupling to SNV distances

Variations in the composition of core genomic content between subspecies of the same species were compared using Jaccard distances. Alongside this, we also calculated sample-pairwise Jaccard distances of detected core gene cluster and Pfam content and compared distances from samples assigned to the same and different subspecies through Wilcoxon signed-rank test.

To determine whether divergence in gene-cluster and Pfam-family content is coupled to nucleotide divergence, pairwise Jaccard distances between subspecies were compared against pairwise SNV distances. As raw SNV and Jaccard distances each combine within-species signal with between-species differences in baseline divergence rate, this relationship was modelled using mixed-effects models with a random intercept for species, fitted independently for gene-cluster and Pfam-family content. The functional form of the relationship was determined by comparing linear, quadratic, and additive (spline-based) mixed-effects models: linear and quadratic models were fitted using the lme4 package, and the additive model was fitted using a generalised additive model (*mgcv::gam*), with the species-level intercept incorporated as a random-effect smooth term and SNV distance modelled with a thin-plate spline (k = 5). Model fit was compared via Akaike Information Criterion (AIC), and, for the nested linear-versus-quadratic comparison, a likelihood-ratio test was also performed, alongside an assessment of the adequacy of the smooth term’s basis dimension using *mgcv::gam.check*. The additive model was selected as the final model based on its improved fit relative to the linear and quadratic alternatives and its ability to capture a non-linear increase in Jaccard distance with SNV distance that plateaus at higher divergence. As a small number of subspecies pairs at the extreme upper end of the SNV-distance distribution disproportionately influenced the fitted smooth, the top 1% of distance values (per gene-cluster/Pfam-family analysis) were excluded prior to final model fitting. A sensitivity analysis confirmed that this exclusion did not qualitatively alter the shape of the fitted relationship, and was necessary to avoid extrapolating an artefactual decline at the sparsely-sampled tail of the distribution (Supp. Fig. S4). Marginal (variance explained by the SNV-distance smooth term alone) and conditional (variance explained by the SNV-distance smooth term together with species identity) R² values were calculated from the corresponding fixed- and random-effect variance components, following the method introduced by Nakagawa & Schielzeth^75^.

### Ribosomal protein codon usage patterns in case study species

To explore additional patterns of divergence within the two species provided as case studies, we calculated the codon frequencies of a set of ribosomal protein genes within each metagenome sample. The codon frequencies were generated using Anvi’o as for the SNV frequencies using ‘--engine CDN’ (see *SNV calling and variability profiles*). The per-sample codon frequencies were subsequently subtracted from the mean codon frequency observed across all samples of the species to derive a per-sample codon usage deviation from the mean. Sample-pairwise distances (Euclidean) were calculated from these codon usage patterns and visualised through a Principal Component Analysis. Codon usage-based separation into subspecies was further assessed through pairwise comparison of Silhouette widths.

## CODE AND DATA AVAILABILITY

The scripts, along with a detailed description of the workflow, to reproduce the analysis presented in this study are available on a public repository: https://github.com/SushiLab/ocean-subspecies-paper/. The data tables containing the information on species and subspecies and that are required by the scripts are available on Zenodo: https://zenodo.org/records/22705862

## ACKNOWLEDGEMENTS

This research was supported by the Swiss National Science Foundation (SNSF) (project grant nos. 205321_184955, 205320_215395), the Swiss State Secretariat for Education, Research and Innovation (SERl) under contract #22.00255 and the European Union (GA#101059915 - BIOcean5D). Views and opinions expressed are however those of the author(s) only and do not necessarily reflect those of the European Union. Neither the European Union nor the granting authority can be held responsible for them. Additionally, Taylor Priest acknowledges funding from the NOMIS Foundation through the NOMIS-ETH Postdoctoral Fellowship program. We would also like to extend our thanks to the ETH IT services and HPC facilities for granting access to the EULER high-performance cluster.

## AUTHOR CONTRIBUTIONS

Key: Concept and Design (CD), Analysis/processing of data (A), Code development (C), Interpretation of results (I), Writing the manuscript (W), Revising the manuscript (R).

Lucas Paoli (CD, A, I, W, R), Taylor Priest (CD, A, I, W, R), Adrián A. Davín (A, I), Melanie L. Stäubli (A, I), Dominic Eriksson (A), Hans-Joachim Ruscheweyh (A), Martin F. Polz (I, R), Shinichi Sunagawa (I, R).

