## Supplementary Information for "Genomically and ecologically cohesive subspecies as biologically meaningful units in the global ocean microbiome"

***Case study 1: Arctic96AD-7 sp002082305, SAR324 Clade***

Based on the union of KEGG orthologs encoded across all three subspecies, *Arctic96AD-7 sp002082305* possesses a versatile metabolic repertoire. It encodes the core enzymatic machinery for carbon fixation via the Calvin-Benson-Bassham cycle (RuBisCo and phosphoribulokinase), alongside a broader complement of genes for heterotrophic growth on organic carbon, including multiple carbohydrate-targeting glycoside hydrolases, long-chain acyl-CoA synthetase for fatty acid activation and multiple sugar ABC transporters. Genes for utilisation of organic sulfur compounds were also evident, including taurine-pyruvate aminotransferase, as well as partial sulfur oxidation machinery (*soxBYZ*). Nitrogen appears to be acquired both inorganically, via an ammonium transporter (*amt* family), and organically, through general amino acid and polyamine (spermidine/putrescine) transport systems. Alongside these metabolic features, the species also encodes a complete flagellar apparatus and chemotaxis system, indicating motility. Together, these metabolic features position this species as a motile, metabolically versatile mixotroph or facultative chemolithoautotroph.

*Arctic96AD-7 sp002082305* was detected across 90 samples spanning all major ocean basins and latitudinal zones. Its three subspecies exhibited distinct biogeographic structuring, which was significantly associated with depth, separating a deep-water subspecies (Subspecies_0) from two epipelagic subspecies (LogOdds ratio 160.53). The epipelagic subspecies were further structured by latitude, with Subspecies_0 restricted to polar waters and Subspecies_2 to tropical and temperate waters (Fig. 5).

Coupled to their occupation of different environmental contexts, the subspecies exhibited distinct signatures of genomic divergence, spanning nucleotide, codon and functional levels. Across all levels, Subspecies_0 and Subspecies_2 consistently emerged as more similar to one another than either was to Subspecies_1. Subspecies_1 also encoded the broadest functional repertoire, with 1,387 unique Pfam families, versus 1,323 in Subspecies_2 and 1,315 in Subspecies_0.

This divergence was further mirrored by subspecies-specific gene content. Subspecies_0uniquely encoded a cyanophycin synthesis/degradation system (*cphA*/K03802, *cphB*/K13282), a mercuric ion transport protein (*merC*/K19058), arsenite transporter (*arsB*/K03325) and a urea-proton symporter (*DUR3*/K20989), which would support enhanced trace-metal handling and nitrogen scavenging in the heterogeneous nutrient availability of the deep oceans. In contrast, Subspecies_1 encoded an enriched repertoire of carbohydrate degradation machinery, including for lyxose (*lyxA*/K09988, *lyxD*/K25260), xylose (*xylA*/K01805), and pentose sugar acid degradation (*araD*/K13876), and uniquely encoded a sensory rhodopsin (*sop*/K04643) and an ECF sigma factor (rpoE/K03088). Subspecies_2 uniquely encoded a near-complete dissimilatory sulfate reduction pathway (*dsrAB*/K11180-1 and *dsrMKOP*/K27187-K27191) alongside additional genes involved in amino acid and aromatic-compound metabolism.

Subspecies-specific signatures of nucleotide-level selection further reinforced this divergence. Subspecies_0 showed elevated pN/pS on arginine/ornithine transport (*aotJ*/K10022: 3.13, versus 0.11 and 0.72 in Subspecies_1 and Subspecies_2), methionine salvage (*mtnA*/K08963: 1.13, versus 0.11 and 0.107), and flagellar biosynthesis genes (*fliO*/*fliZ*/K02418: 3.83, and *fliW/*K13626: 2.46)*,* alongside purifying selection on heat shock protein *hspQ* (K11940: 0.0015 versus 0.15 and 0.075) and translation initiation factor IF-1 (K02518: 0.0015, versus 0.038 and 0.031). Beyond these, subspecies-specific selective signatures were also identified across 53 additional genes in the shared genomic backbone, spanning diverse functional categories including transport, cell division, translation, and stress response, with no strong convergence on a single pathway or process.

***Case study 1: AAA300-D14 sp002715065, Gammaproteobacteria***

The genomic repertoire of AAA300-D14 sp002715065 points to an aerobic, heterotrophic lifestyle built around the degradation of complex organic matter. Multiple glycoside hydrolases suggest a capacity to use polysaccharides as a carbon source, complemented by a complete TCA cycle and an aa3-type cytochrome c oxidase, indicative of aerobic respiratory metabolism. Nitrogen appears to be acquired primarily through exogenous ammonium, via an amt-family transporter. The present of a complete flagellar apparatus suggests a motile capacity. The presence of a TonB-dependent iron-acquisition system provides the baseline capacity for iron uptake, which is further supplemented by subspecies-specific iron specialisation, structured along an iron-availability gradient across the species' tropical Atlantic, Pacific and Indian Ocean distribution.

Subspecies within AAA300-D14 sp002715065 diverged only modestly in gene content. Subspecies_0 and Subspecies_1 were more similar to one another (Jaccard distances of 0.221 for gene clusters, and 0.045 for Pfam families) than either was to Subspecies 2 (0.363 and 0.081) (Fig. 6c), and the differences in functional repertoire size across the subspecies ranged from 1,110 (Subspecies_2) to 1,200 (Subspecies_1) unique Pfam families. This divergence was overwhelmingly concentrated in iron-acquisition machinery: Subspecies_1 uniquely encoded a heme substrate-binding protein and permease (*hmuT*/K25132, *hmuU*/K25133), an iron-complex ABC transporter ATP-binding domain (*ABC.FEV.A*/K02013), and a Fe(3+)-dicitrate transporter (*fecA*/K16091), alongside the highest copy number of the TonB-associated *exbD* components. Similarly, Subspecies_2 harboured the highest copy number of the iron complex outer membrane receptor (*fhuA*/K02014) and the TonB-associated *exbB* components. Beyond this, subspecies-specific gene content was otherwise limited. Subspecies_0's unique content, encompassing a formate dehydrogenase subunit, one flagellar regulatory genes, and a enol-CoA isomerase enzyme, spans unrelated functions with no shared theme, while Subspecies_2 encodes only a single uniquely annotated KEGG ortholog (K15770), consistent with its more functionally streamlined gene-cluster content, although this perspective may, in part, reflect incomplete KEGG annotation coverage.

We identified 128 species' core genes that exhibited subspecies-specific pN/pS signatures. Four of these were associated with iron metabolism: Flavodoxin II (*fldB*/K03840), which was under strong purifying selection in Subspecies_1 but positive selection in Subspecies_0 and

Subspecies_2,; iron-sulfur cluster assembly protein cyaY (K06202) and the cytochrome c maturation protein ccmE (K02197), which both showed subspecies-specific differences in purifying selection pressure, particularly across Subspecies_0 and _1; and the ubiquinol-cytochrome c oxidase iron-sulfur subunit (K00411) that also showed pronounced purifying selection in Subspecies_0 (pN/pS = 0.003, versus 0.062 and 0.039). Beyond these, two independent genes in the ubiquinone (coenzyme Q) biosynthesis pathway stood out: chorismate lyase (*ubiC*/K03181) showed markedly elevated pN/pS in Subspecies_1 (1.46, versus 0.26-0.31 in the other two subspecies), and the pathway's accessory factor (*ubiJ*/K03690) showed a comparable pattern in Subspecies_2 (1.84, versus 0.25-0.27). The recurrence of this signal across two independent genes in the same pathway, albeit in different subspecies, suggests that ubiquinone biosynthesis may be a recurring target of relaxed selective constraint in this species.

### Supplementary Figures

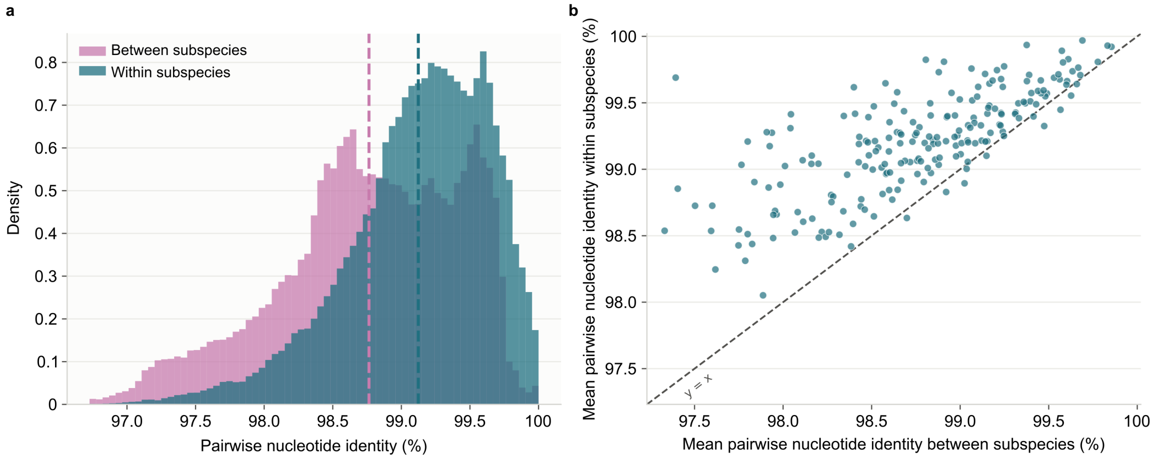


**Supplementary Figure S1. Subspecies divergence on the species' core genomic backbone.** For each species, we determined the set of single-copy marker genes as the Clusters of Orthologous Groups (COGs) that were identified across all species' samples (after applying the ≥90% species core genome coverage threshold). Each core gene was aligned across samples using MAFFT and subsequently concatenated into a per-species nucleotide supermatrix. Pairwise nucleotide identity between metagenome samples was defined as the proportion of matching positions over all sites at which both samples carried an unambiguous, non-gap base (pairwise deletion). **a)** Pairwise nucleotide identity of metagenome samples assigned to the same (within) subspecies or different (between) subspecies within the same species. Dotted lines represent mean values. **b)** Mean pairwise nucleotide identity of the within versus between subspecies comparisons within each species. Each dot represents one species.

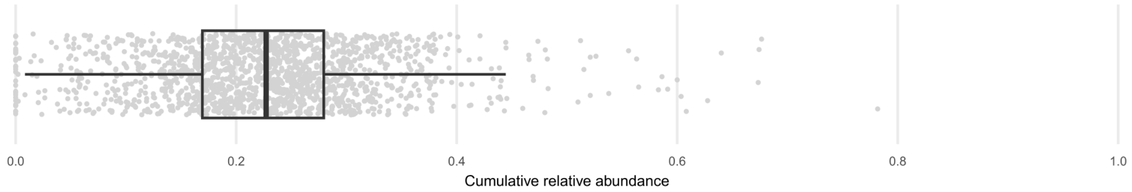


**Supplementary Figure S2. Abundance of species across OMD.** The 270 species with subspecies detected in this study account for an average of 23% of the relative abundance of microbial communities across the 1’669 metagenomes as estimated by mOTUs profiles.

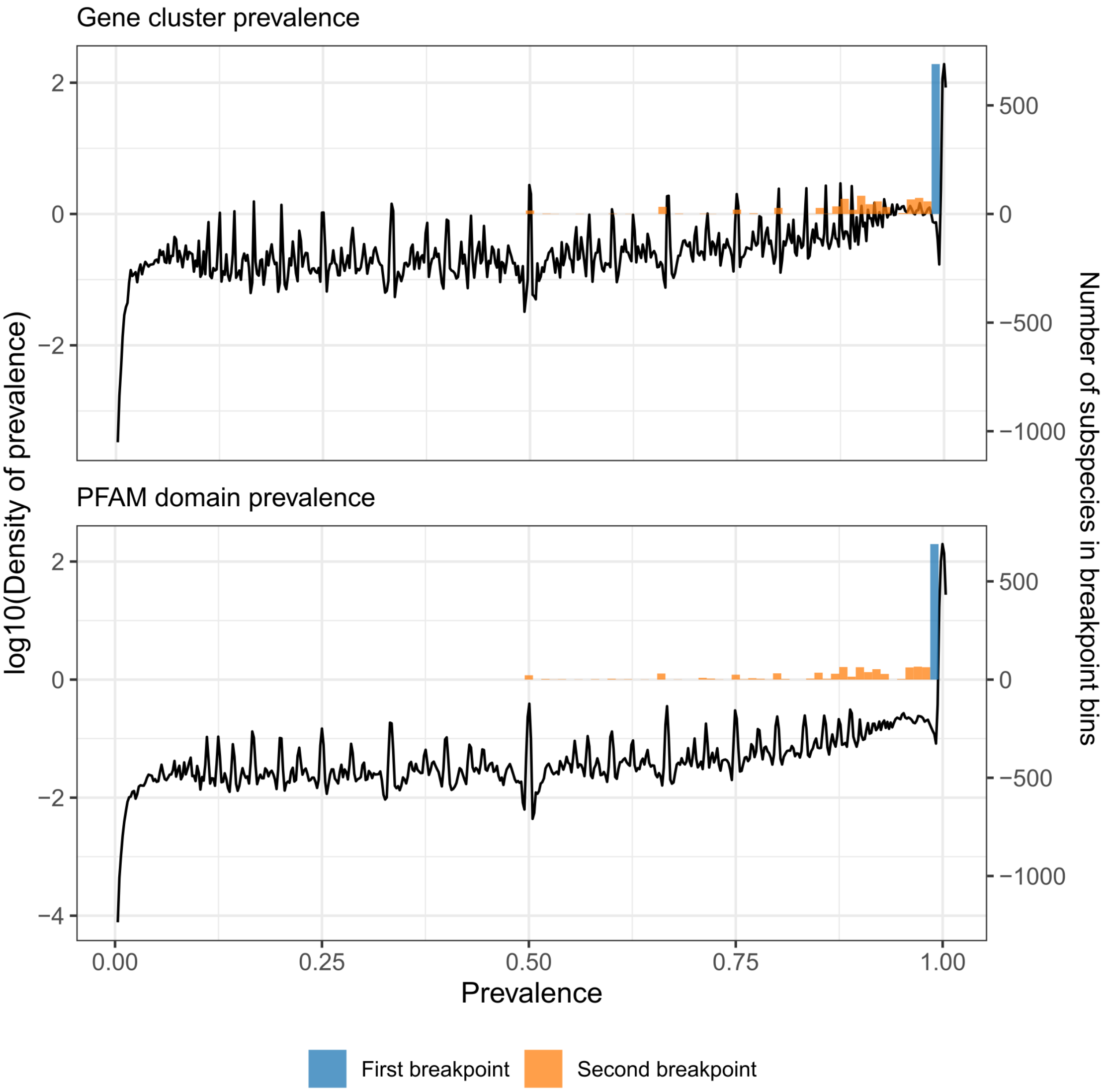


**Supplementary Figure S3. Distribution of gene prevalences across species and subspecies samples.** To determine the core genome of each subspecies, we assessed the prevalence of detected gene clusters and Pfam families across samples. Prevalence values were subsequently grouped into 1% bins and the two primary inflection points (in a descending order from 100%) were determined. The lines represent the density of observations across the 1% prevalence bins, while the bars represent the inflection points, with their heights capturing the number of species or subspecies that shared the point.

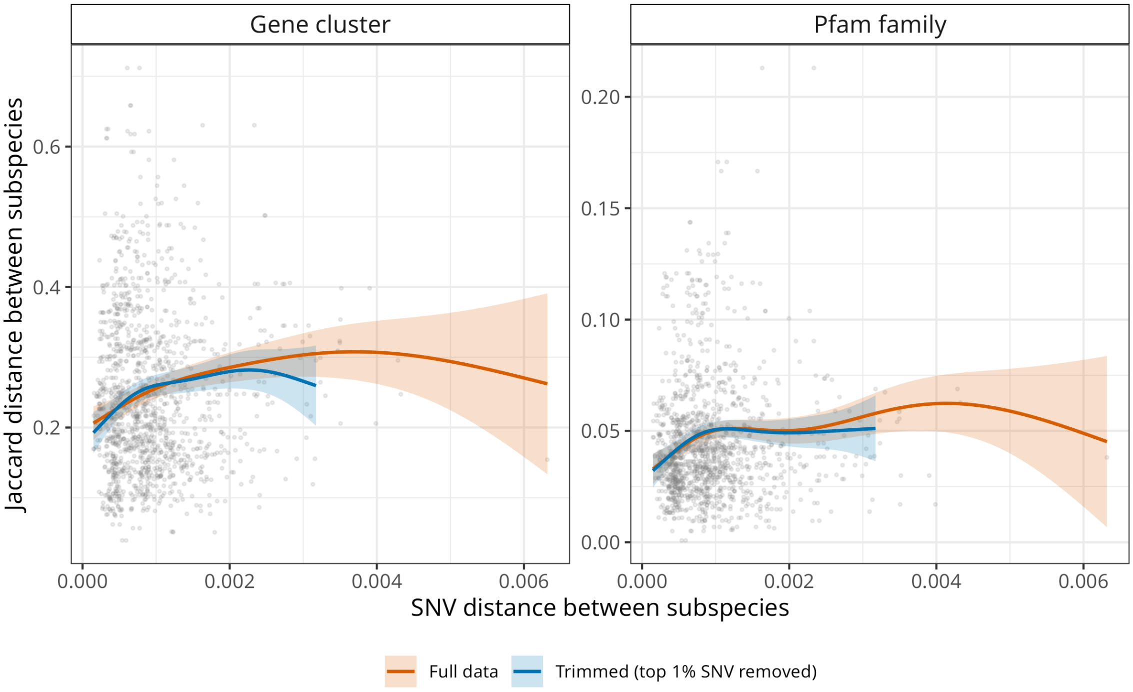


**Supplementary Figure S4. Sensitivity of the GAM-fitted relationship between SNV distance and gene-content divergence to exclusion of the sparsely-sampled upper tail.** Generalised additive model (GAM) smooth fits describing the relationship between pairwise SNV distance and Jaccard distance in core gene-cluster content (left) and core Pfam-family content (right) between subspecies of the same species. Points show individual subspecies pairs. Orange lines and shading show the fitted smooth and 95% confidence interval using the full dataset; blue lines and shading show the corresponding fit after excluding the top 1% of SNV-distance values per analysis, which was used for the final model presented in the main text (Fig. 3e).

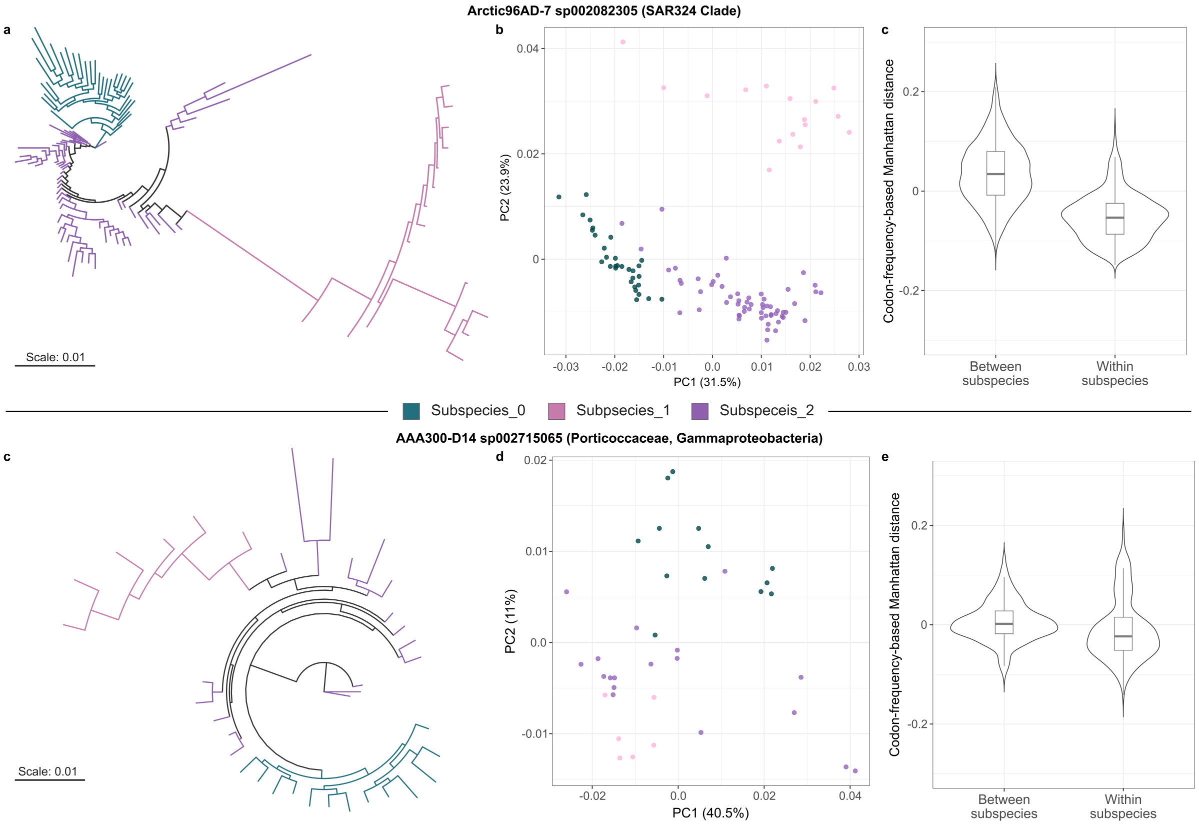


**Supplementary Figure S5. Genomic signals of divergence within the two case study species. a)** and **c)** Phylogenetic reconstruction of the species' core genome. For each species, we extracted the consensus sequence of each gene in each metagenome sample from the SNV profiles and subsequently aligned sequences using MAFFT. Individual gene alignments were concatenated into a supermatrix representing the whole species core genome and a maximum likelihood phylogenetic tree was reconstructed using IQTREE using the best-fit substitution model selected by ModelFinder (parameters: -m MFP), with 1000 ultrafast bootstrap replicates (parameters: -B 1000). Leaves represent metagenome samples while terminal branches are coloured according to the subspecies assignments of each sample. **b, c, d, e)** Comparison of the codon usage bias within ribosomal proteins across samples. Codon usage frequencies in ribosomal proteins were calculated for each metagenome sample. Per-sample codon frequencies were subsequently subtracted from the mean codon frequency observed across all samples of the species to derive a per-sample codon usage deviation from the mean. Sample-pairwise distances (Euclidean) were then calculated from these codon usage deviations and visualised through a Principal Component Analysis (**b** and **d**) - wherein each point represents a metagenome sample coloured according to its subspecies assignment. Pairwise distances were also grouped according to whether they represent within- or between-subspecies comparisons and visualised as violin and boxplots (**c** and **e**) - wherein boxplots capture the interquartile range and the median, with whiskers representing the 1 x standard deviation of the interquartile range.

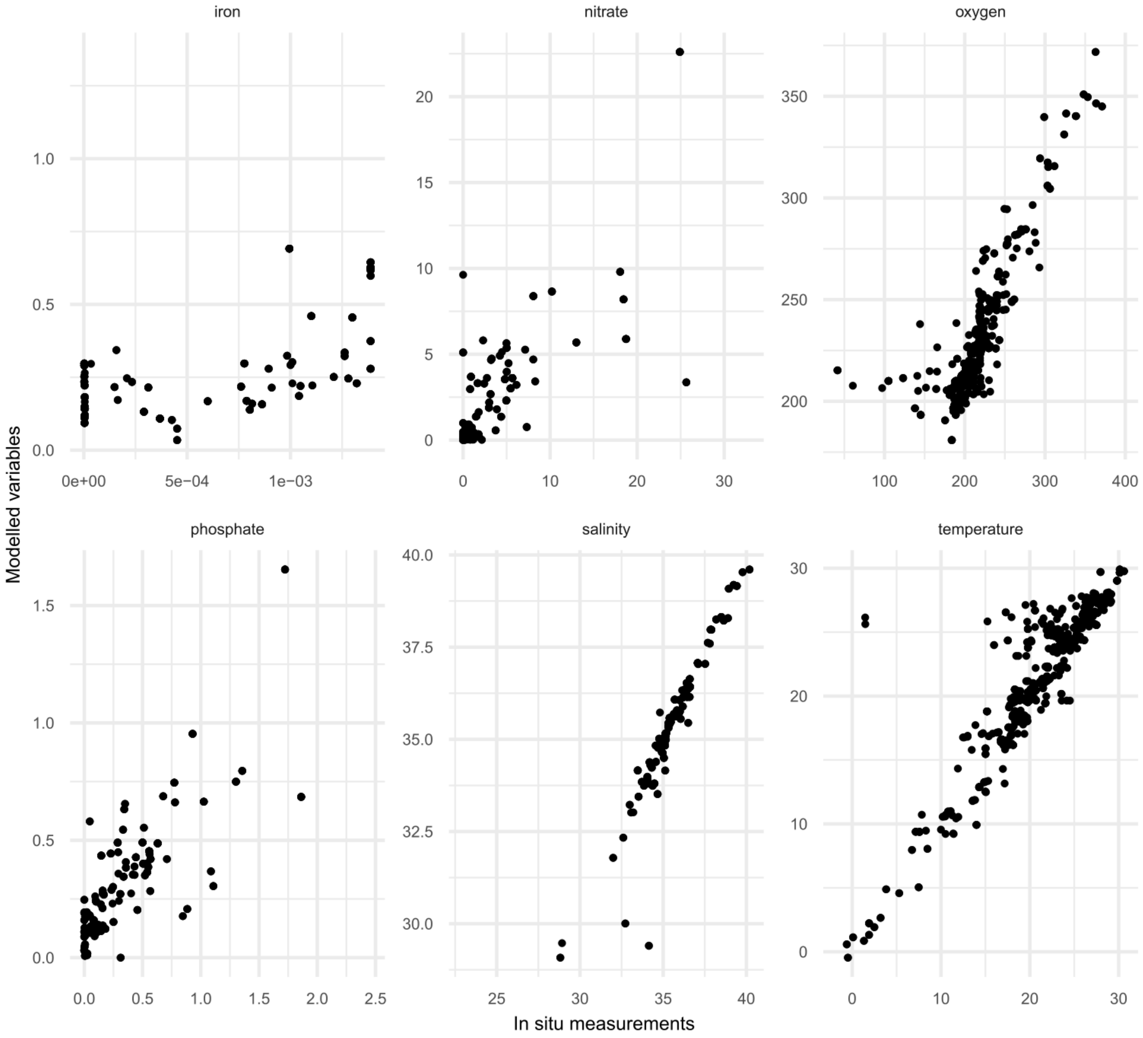


**Supplementary Figure S6. Comparison of in situ and modelled environmental parameters.** Correlations between modelled environmental parameters and in situ measurements in the first 100m of the water column.
